# Regulatory Bias Constrains Epigenetic Aging Trajectories

**DOI:** 10.64898/2026.07.22.740004

**Authors:** Taisiia Morozova, Marina Axelson-Fisk, Annikka Polster

**Affiliations:** Department of Life Sciences, Chalmers University of Technology, Gothenburg, Sweden

**Author notes:** Contributing authors. These authors contributed equally to this work (Marina Axelson-Fisk, Annikka Polster).

**Keywords:** Epigenetic aging, DNA methylation, Stochastic processes, regulatory bias, stochastic modeling, simulation

## Abstract

Aging reflects both stochastic fluctuation and biological regulation. We present a Markov chain framework for epigenetic aging that extends noise-driven models by adding a state-dependent bias term representing regulatory constraint. Using DNA methylation data from mice, rats, and bats, we show that empirical epigenetic aging is characterized by a progressive restriction of the accessible state space. We demonstrate that a stochastic model incorporating state-dependent regulatory bias successfully reproduces this constraint, whereas standard noise-driven models fail to capture it. Subsequently, the results indicate that a regression-based drift model can be used to predict future trajectories and achieve lower mean, covariance, and state-increment dependence errors than the biased model. The pattern of state loss is consistent with discrete bifurcation events, suggesting resilience declines stepwise rather than continuously. This implies that the solution space for intervention narrows irreversibly at each transition, making intervention timing critical.

## 1 Background

Aging is a complex biological process characterized by a gradual decline in physiological function and an increased risk of disease and death. At the molecular and cellular levels, aging involves a wide range of interconnected factors, including genomic and epigenetic changes and proteostasis loss. Over the past decades, extensive research has led to a wide range of theories and perspectives on aging, with researchers seeking to approach aging from various angles. The most common ways of representing aging are molecular damage accumulation, trade-offs between reproduction and maintenance, and the gradual loss of regulatory control. Although the precise definition of aging remains an open challenge, the existing research provides the key insight: aging is not driven by a single mechanism, but rather emerges from the interaction of many processes across different biological scales [1, 2]. As a result, aging should be viewed as a dynamic process of complex interactions between stochastic effects and regulatory constraints, rather than a constant, gradual decline.

In recent years, *aging clocks* have become powerful tools for quantifying biological age [3, 4]. These models, often based on high-dimensional data such as DNA methylation, gene expression, or proteomics, can predict biological age with remarkable accuracy. However, this predictive ability comes with imitations. While such models can estimate *how old* a system appears biologically, they generally do not explain *why* the system has reached that state. For instance, an increase in predicted biological age may reflect changes in multiple genes or pathways, but standard models do not identify which factors are driving these changes or how they interact over time. This leads to a broader challenge in aging research: the gap between prediction and understanding. High-dimensional models can summarize complex biological states into a single number; however, this number is not enough to fully understand the aging process. This motivates the need for frameworks that not only describe the current state of a system but also explain how it evolves and most importantly, which paths are affected the most.

From a biological perspective, aging is inherently dynamic and subject to variability across individuals and even across cells within the same organism. This variability arises from a combination of factors: molecular fluctuations, environmental influences, and regulatory mechanisms. As a result, it is natural to model aging as a stochastic process, where each state of the system evolves according to a *state-dependent probability distribution*. In this framework, the state at a given time point represents a high-dimensional biological profile, such as gene expression or epigenetic markers, while transitions between states capture the accumulation of molecular changes. A further simplification is to assume that the process is Markovian, meaning that the current state contains all relevant information about the past. Biologically, this is justified by the fact that molecular states already encode all the cumulative effects of prior changes, so that the explicit history of previous states becomes non-informative once the present configuration is known. Although aging can indeed me modeled as such a process, it is not purely random and its dynamics are also structured by regulatory constraints, feedback mechanisms, and environmental pressures, as indicated by the recent studies [5]. These factors influence how states evolve, making certain transitions more or less likely than others, and some even impossible after having reached a certain state. A concrete example of such constrained and partially irreversible dynamics can be found in neurodegenerative diseases such as Parkinson’s disease. Molecular changes involving genes such as *SNCA* and *LRRK2* drive the system toward pathological states, in particular through the aggregation of *α*-synuclein and Lewy-body formation [6, 7]. Once critical thresholds are crossed, such as sustained protein aggregation or mitochondrial dysfunction, recovery to a previous healthy state becomes highly unlikely[8]. As a result, the system evolves within a restricted region of the state space, where transitions back to earlier states are inaccessible.

This illustrates that biological dynamics of aging are not only noisy but also exhibit trends and path dependence. Hence, purely random evolution is not sufficient to it. We therefore propose to define aging as a time series model that captures both random and structural effects. The idea is that, at each time point in life, the state of the organism evolves based on the combination of three terms: the state at the previous time point, random noise, and an additional state-dependent term. We refer to the latter as *regulatory bias*, which reflects how biological mechanisms shape and constrain possible state trajectories as the organism ages. This perspective is supported by recent work in single-cell biology and dynamical modeling. For example, trajectory inference methods aim to reconstruct the directions of biological systems evolving over time from snapshot data [9, 10], while approaches such as RNA velocity estimate short-term changes in gene expression and show a notion of direction in the state space [11]. Generally, aging is increasingly modeled as a stochastic process evolving in a high-dimensional state space, where individuals transition through a sequence of physiological and molecular states over time [3, 12, 13]. These stochastic approaches are particularly well-suited to aging research because longitudinal data are often sparse, noisy, and incomplete. DNA methylation aging provides a particularly relevant example of this framework. Age-related methylation changes are often described using stochastic models of epigenetic drift, in which random methylation gains and losses accumulate throughout life [14, 15]. While such models capture many observed features of aging and form the basis of epigenetic age predictors, they do not explicitly account for mechanisms that may bias transitions between methylation states. The nature of such bias is unlikely to be universal and may depend on tissue type, genomic context, and species-specific regulatory programs ([14]). However, one consequence of such dynamics would be a progressive reduction in state-space exploration, causing trajectories to become increasingly confined to a smaller set of methylation states.

In this paper, we focus on modeling epigenetic aging and hypothesize that aging is accompanied by a progressive reduction in the accessible methylation state space. We further hypothesize that this contraction is driven by a form of regulatory bias that alters the probabilities of transitions between methylation states, making some trajectories more likely than others. To capture this effect, we extend a stochastic framework by incorporating a bias term, allowing biological regulation to influence the evolution of DNA methylation patterns with aging. Under this interpretation, aging is not solely characterized by stochastic fluctuations but also by regulatory constraints that progressively limit the range of accessible methylation states. Moreover, this framework connects naturally with statistical and machine learning methods. Using the observed methylation data, transition dynamics can be estimated directly through regression-based models, enabling the full characterization of methylation-state evolution and the prediction of previously unobserved future states.

### 1.1 Our contribution

To capture these ideas, we model epigenetic aging as a time-inhomogeneous Markov process with an additional bias (drift) term evolving over time. At each time point (t), the state *X*_*t*_ *∈* ℝ^*d*^ represents a (d)-dimensional vector of DNA methylation levels across selected CpG sites. Time-inhomogeneity implies that transition probabilities change with age, so that the likelihood of moving between methylation states depends both on the current state and on time. We then conduct a series of experiments to investigate how regulatory bias influences the range of methylation states accessible during aging. First, we investigate the role of regulatory bias by simulating multiple stochastic processes starting from the same initial state derived from real data. These include an unbiased process driven solely by random noise and several biased processes incorporating different forms of state-dependent drift. We compare these trajectories with the empirical data in terms of states reachability, exploration, and variability, and show that all biased processes reproduce the progressive loss of the accessible methylation states observed in the data, whereas the unbiased process does not. We then estimate the transition dynamics directly from data using biased and unbiased approaches. At each time step, we learn the transition distribution using regression-based methods and generate future states accordingly. We then evaluate predictive performance against observed trajectories and demonstrate that the model incorporating regulatory bias consistently provides a more accurate representation of the aging dynamics.

## 2 Effect of regulatory bias

The initial goal of this paper is to investigate whether epigenetic aging is accompanied by a progressive restriction of the accessible state space and whether this behavior is better reproduced by a stochastic model with state-dependent regulatory bias than by a purely noise-driven model. We show that empirical methylation dynamics become increasingly constrained with age and that the biased model captures this behavior more accurately than the unbiased model. We therefore begin with an extensive simulation study to examine how the presence of regulatory bias influences the evolution of the system. The idea is to simulate two processes with the same initial state: one is purely the accumulation of stochastic noise, and the other includes an additional state-dependent bias term. The former and the latter are referred to as unbiased and biased processes, respectively. In the subsequent experiments, the model will be initialized using real methylation data from mice, rats, and bats, which will then be used for validation. From this data, forward simulations will be generated. The resulting simulated trajectories will be evaluated using a range of quantitative metrics, including state reachability, exploration rate, and repeat rate, with the aim of identifying potential reductions in the set of accessible states induced by the bias. For validation, these simulated trajectories will be systematically compared to empirical time series derived from the validation dataset. This allows us not only to demonstrate that the presence of a regulatory bias alters stochastic outcomes, but also to provide evidence that such a bias indeed exists in empirical data.

### 2.1 Data description

To validate the proposed framework, we use publicly available DNA methylation datasets from the Gene Expression Omnibus (GEO), each containing methylation measurements collected across multiple ages and species. The datasets include mouse, rat, and bat methylation profiles spanning different tissues and age ranges. The first dataset, GSE120132 [16], contains reduced representation bisulfite sequencing (RRBS) methylation data from *Mus musculus* collected across the full lifespan and multiple tissues, with the mice ages ranging from 2 to 20 months. From this dataset, we restrict the analysis to kidney tissue samples only. The second dataset, GSE164127 [17], contains mammalian methylation array data from multiple bat species and tissues. We select the species *Phyllostomus hastatus* (Greater spear-nosed bat), with 115 observations aged 1–19 months). The third dataset, GSE161141 [18], contains RRBS methylation measurements from Fischer 344 rats spanning ages 1–27 months; all available rat samples are included in the analysis. Finally, GSE80672 [19] contains mouse whole-blood RRBS methylation profiles for ages 0.5–35 months and experimental conditions, from which all 255 samples are used.

For each dataset, methylation beta values are organized into a three-dimensional tensor of size *time × observations × features*. Each observation is represented as a vector *X*_*t*_ *∈* ℝ^*d*^ containing methylation levels for *d* CpG sites at a given age. To obtain a balanced temporal representation, samples were uniformly subsampled within each age group, yielding 3–7 observations per time point, while samples with missing age information were excluded. To address the high dimensionality of the methylation data, analyses were performed on randomly selected subsets of *k* CpG sites. For each value of *k*, estimates were computed across *n* = 100 bootstrap samples and averaged. This procedure additionally allowed us to assess the robustness of the results with respect to the number of selected CpG sites. To avoid degenerate feature subsets, only bootstrap samples for which at least 20% of the selected CpG sites exhibited positive variance across observations at every time point were kept.

### 2.2 Stochastic model

We investigate the effect of regulatory bias by simulating and comparing two stochastic processes in the context of aging dynamics inferred from DNA methylation data. Specifically, we model epigenetic aging through the temporal evolution of methylation profiles and compare (i) an unbiased process driven purely by stochastic fluctuations and (ii) a biased process incorporating an additional state-dependent drift term.

The stochastic processes are initialized using DNA methylation data from several independent datasets described in Section 2.1. Each experiment is conducted separately for a given dataset, so that both the biased and unbiased processes are initialized exclusively from that dataset. The processes are initialized using the observed methylation states at the earliest available time point (for instance, in the case of GSE132741, the youngest species is measured at the age of 2 months, hence *t*_0_ = 2), after which the subsequent methylation trajectories are simulated. The data are represented as a three-dimensional object indexed by time, observations, and features. Let *T* denote the total number of time points, *n* the number of observations (individuals), and *p* the number of methylation features (CpG sites). Since methylation profiles are expressed through beta values, all data is limited to the interval [0, 1]. Hence, at each time point *t* = 0, …, *T*, the state of the system is represented by a matrix

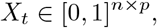

where each row corresponds to one individual and each column corresponds to the same methylation feature of different individuals. Thus, the *i*-th row of *X*_*t*_ contains the methylation vector for individual *i* at time *t*. Note that observations are independent, and hence each row can be interpreted as an independent multi-dimensional stochastic process whose state space consists of *p*-dimensional methylation vectors. For convenience, we represent all observations jointly as a matrix-valued process. This matrix formulation also allows us to later analyze and learn the collective transitions dynamics of the methylation state space.

As a base model, we use a multi-dimensional time-homogeneous Markov chain driven by normal noise only. Since methylation values are naturally bounded between 0 and 1, each state is sampled from a multivariate normal distribution truncated to [0, 1]^*d*^ as

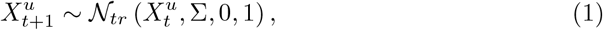

where Σ is a fixed covariance matrix. Here, *N*_*tr*_(*µ*, Σ, 0, 1) denotes a multivariate normal distribution with mean *µ* and covariance Σ, truncated to the interval [0, 1] in each coordinate. Equivalently, samples are drawn from *N*(*µ*, Σ) and retained only if all components lie within the biologically meaningful methylation range [0, 1]. The last two arguments specify the lower and upper truncation bounds, respectively. The covariance matrix Σ controls both the magnitude of the fluctuations and the dependence structure between CpG sites.

### 2.3 Bias formulation

The process (1) has been proposed to model biological aging in the studies of aging clocks [3]. This process represents pure stochastic diffusion and accumulated noise and, as was shown in [3], can be used for robust prediction of the biological aging. We argue that, although such a representation of aging is sufficient to predict the biological age, it does not efficiently capture some important aging-related effects, such as regulatory homeostasis and coordinated molecular adaptation. Although the process (1) is not entirely unbiased since the truncation at [0, 1] induces an effective drift, we argue that this drift is not an adequate representation of biologically driven age-related changes. Hence, we introduce a more complex model to capture the aging dynamics, which accounts not only for the stochastic diffusion but also for regulatory drifts in the observed values. Biologically, molecular states such as DNA methylation levels or gene-expression activity are additionally driven by regulation [20–23]. Mathematically, such regulation mechanism can be understood as a stochastic process with a restoring drift toward a population-level average value. Consequently, when stochastic changes drive molecular values away from “typical” (average) states, regulatory processes may induce restoring dynamics that partially push these values back toward the mean. To model such dynamics, we refine the process (1) by introducing the additional state-dependent drift term. The biased aging process is then modeled as a *time-inhomogeneous* Markov chain

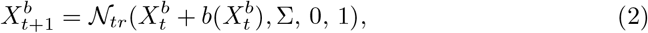

where the bias has the form of

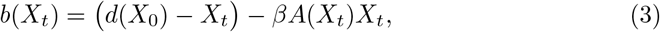

and 0 *< β <* 1 controls the bias strength. Here, *d*(*X*_0_) = (*µ*_1_, …, *µ*_*p*_) is a fixed vector of mean methylation values computed across individuals at the earliest time point. Biologically, it represents a population-level reference state and models regulatory mechanisms that tend to restore methylation levels toward typical values observed in young individuals. The matrix

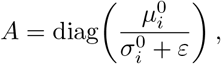

Where 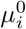 and 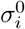 denote the population mean and standard deviation of feature *i* estimated from the empirical data at the earliest available time point. The matrix *A* is computed once prior to simulation and remains fixed throughout the process. If only one observation is available and *X*_*t*_ is a vector, then the diagonal of *A*(*·*) is given component-wise by the absolute value of *X*_*t*_. The first term in (3)

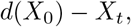

acts as a mean-reverting drift. Features with values below the population mean tend to increase, whereas features above the mean tend to decrease. This term comes from classical auto-regressive models of order 1 (AR(1), [24, Section 3.4]) and its purpose is to push values toward the population mean. However, all genes are differently affected as the organism evolves, which means that we must control not only *where* the gene values move, but also *how strongly* they are affected. For this reason, we strengthen the drift by additionally putting weight on each of the features. The second term in (3)

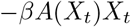

reflects feature-specific regulation. In other words, features with large mean expression and low variability receive a stronger drift, while highly variable features are affected more weakly. Intuitively, the model assumes that genes with stable methylation patterns are more biologically meaningful than genes whose expression changes mostly due to random fluctuations. Subsequently, stable features experience stronger systematic drift, while highly variable features are affected more weakly. To better explain the meaning of the proposed model, we provide a simple numerical example. Consider a small dataset with 2 features and 2 observations, so that at some time point *t* we have

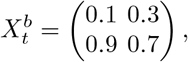

and let

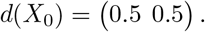

The feature-wise standard deviations compute from the initial data *X*_0_ are

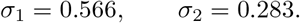

Thus,

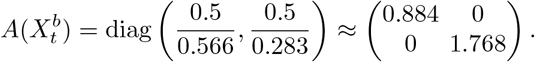

The first feature has larger variability than the second feature, so its coefficient in 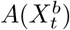 is smaller. Consequently, the second feature is adjusted more strongly. Using the update rule with *β* = 0.3 and ignoring noise gives

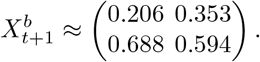

The first row increases because its values are below the feature-wise mean, while the second row decreases because its values are above the mean. At the same time, the second feature changes more strongly due to its lower variability and therefore larger bias coefficient.

### 2.4 Choice of bias term

While the proposed bias formulation in (3) is indeed motivated by biological considerations and feature-specific regulation, the exact form of such bias in methylation dynamics is generally unknown and may vary depending on the data, species, and tissues. Biologically, regulation in genes during aging may be encoded in various ways, and therefore several alternative bias formulations are plausible. To investigate the sensitivity of the aging dynamics to the choice of bias mechanism, we consider several alternatives in addition to the proposed model, which are summarized in Table 1. We briefly explain the biological meaning of the proposed approaches. The FS approach retains the feature-specific weighting matrix *A*(*X*_*t*_) but removes the global restoring force toward the reference state *d*(*X*_0_). This formulation completely removes the contribution of feature-specific regulation and allows us to assess whether the feature-based weighting alone is sufficient to constrain the aging dynamics. The LMR model consists of a purely linear restoring mechanism. Here, all features are pulled toward the population-level state with equal strength. Consequently, the model captures global regulation but ignores feature-specific differences in variability. The NMR approach introduces a nonlinear restoring mechanism through a quadratic term. Unlike the linear model, the strength of regulation increases with the magnitude of the deviation from typical values. As a result, features with large deviations from the mean are penalized more strongly, producing a form of nonlinear damping.

**Table 1:**
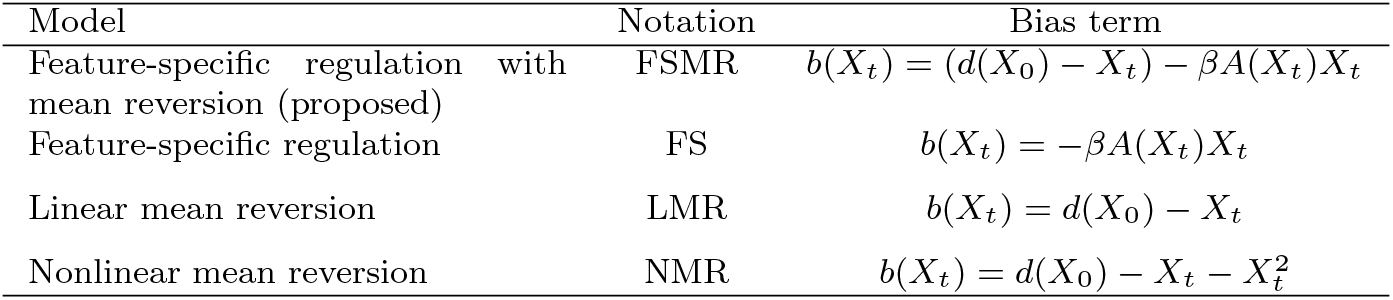
Bias formulations considered in the comparative study.

| Model | Notation | Bias term |
| --- | --- | --- |
| Feature-specific regulation with mean reversion (proposed) | FSMR | $b(X_t) = (d(X_0) - X_t) - \beta A(X_t)X_t$ |
| Feature-specific regulation | FS | $b(X_t) = -\beta A(X_t)X_t$ |
| Linear mean reversion | LMR | $b(X_t) = d(X_0) - X_t$ |
| Nonlinear mean reversion | NMR | $b(X_t) = d(X_0) - X_t - X_t^2$ |

### 2.5 Experiment setting and results

To evaluate the effect of regulatory bias on the dynamics, we conduct a series of simulation experiments, subsequently using data from Section 2.1 for initialization and validation. For each dataset, we use its initial state *X*_0_ to initialize all simulated processes. We then simulate several biased processes (2), each corresponding to a different bias formulation described in Table 1, as well as a reference unbiased process according to (1). To ensure a fair comparison, all simulated trajectories are generated with the same number of features, observations, and time points as the validation dataset. The simulated processes are further assessed using a collection of explorability metrics and compared against the corresponding metrics computed from the real data. First, we measure the *visited-state growth*, defined as the cumulative number of unique methylation states visited by the process over time. Since methylation values can take any value between 0 and 1, two observations are rarely exactly identical. To account for this, we group similar methylation profiles together. For example, suppose two CpG sites have methylation values ((0.23,0.27)) and ((0.26,0.24)) at two different time points. Using bins of width 0.1, both profiles are assigned to the state ((0.2,0.2)). In this case, the methylation state between the two time points has not changed. In contrast, if the second profile is ((0.34,0.28)), it is assigned to the state ((0.3,0.2)), indicating a transition to a different state. The visited-state growth is then the cumulative number of different states observed over time. As a second metric, we compute the *exploration rate*, defined as the number of newly visited states at each time step.

While the visited-state growth captures cumulative behavior, the exploration rate gives us a local view of how quickly the process continues to discover new states. A decreasing exploration rate indicates that the process eventually loses access to certain states and becomes confined to a smaller subset of states. Third, we evaluate the *repeat rate* of the methylation state distribution at each time step. This metric measures how often the process returns to states that have already been visited. A high repeat rate indicates that the process repeatedly explores the same regions of the methylation state space, whereas a low repeat rate suggests continued exploration of new states. All metrics are computed for the real data, the unbiased process, and each of the biased processes described in Table 1, and compared over time. The aim of this experiment is to test whether a stochastic model with state-dependent regulatory bias better reproduces the progressive restriction of the accessible methylation state space observed during epigenetic aging. We therefore expect biased and real processes to have lower visited-state growth and exploration rate than the unbiased one, and, respectively, a higher repeat rate. With this approach, we can assess whether the regulatory bias is indeed observed in real data, and in particular, which of the proposed biased models in Table 1 better capture the exploration dynamics of epigenetic aging. The results are presented in Figures 1–3.

**Fig. 1:**
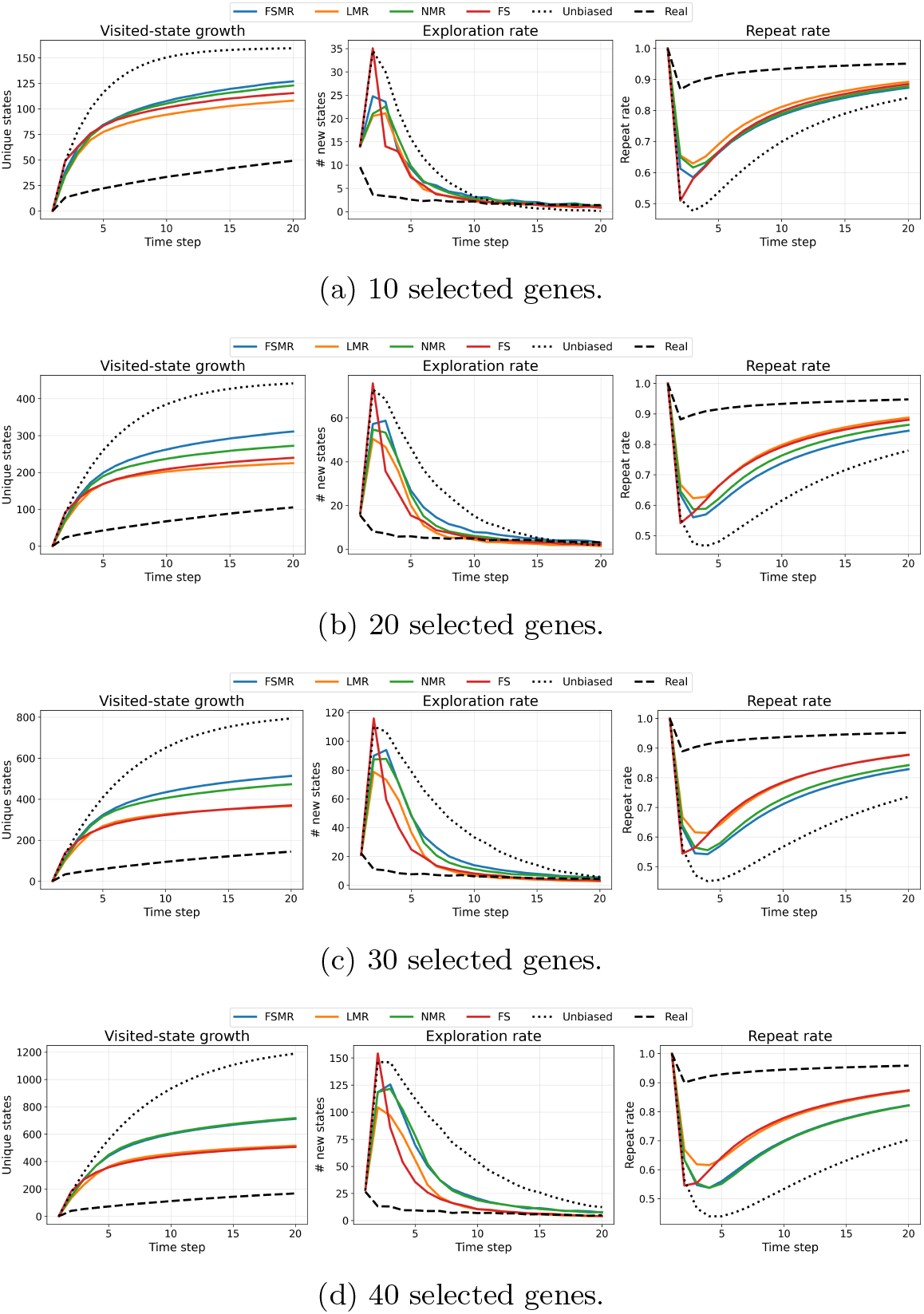
Dynamics comparison for dataset GSE161141 across different feature subset sizes.

**Fig. 2:**
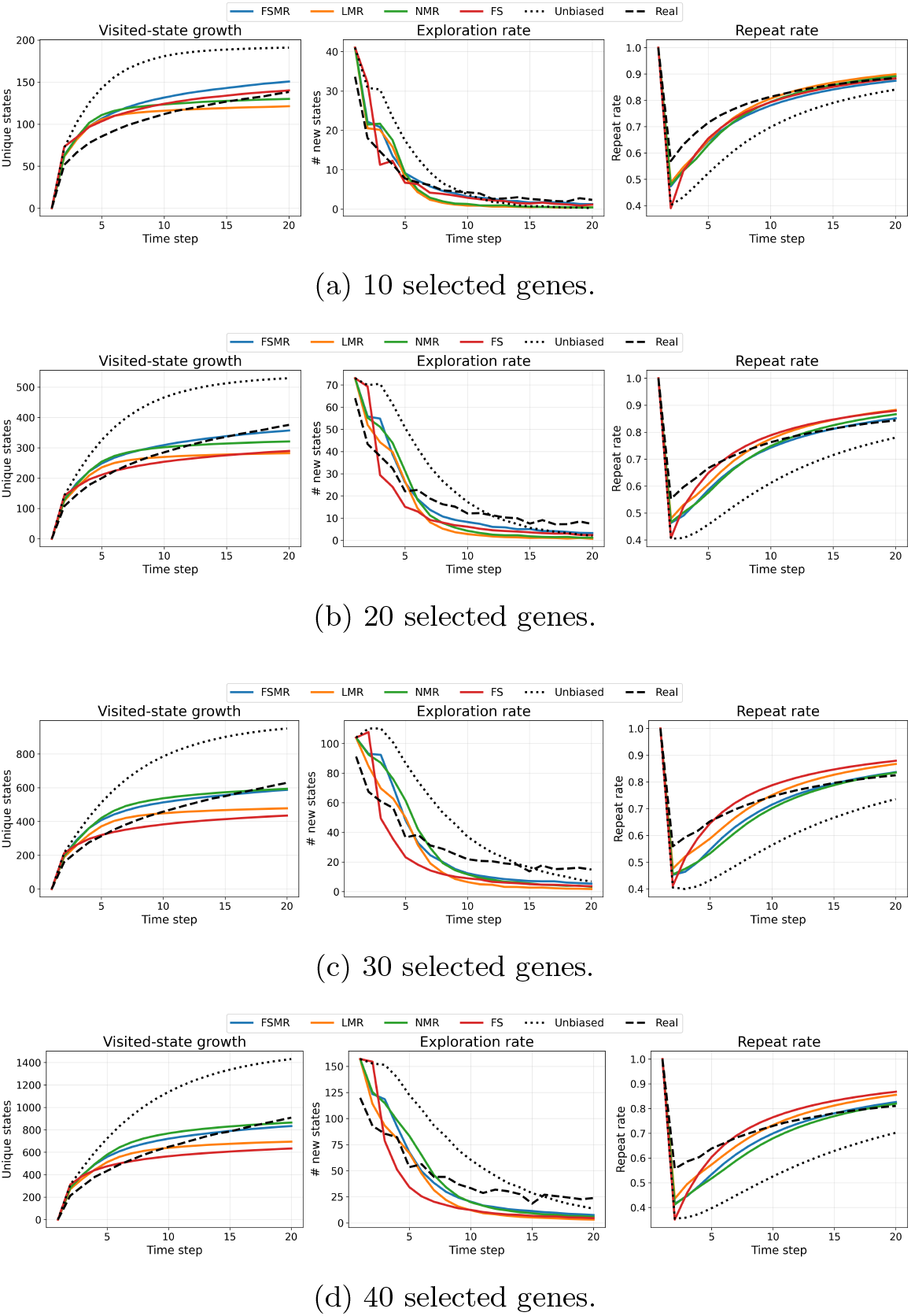
Dynamics comparison for dataset GSE80672 across different feature subset sizes.

**Fig. 3:**
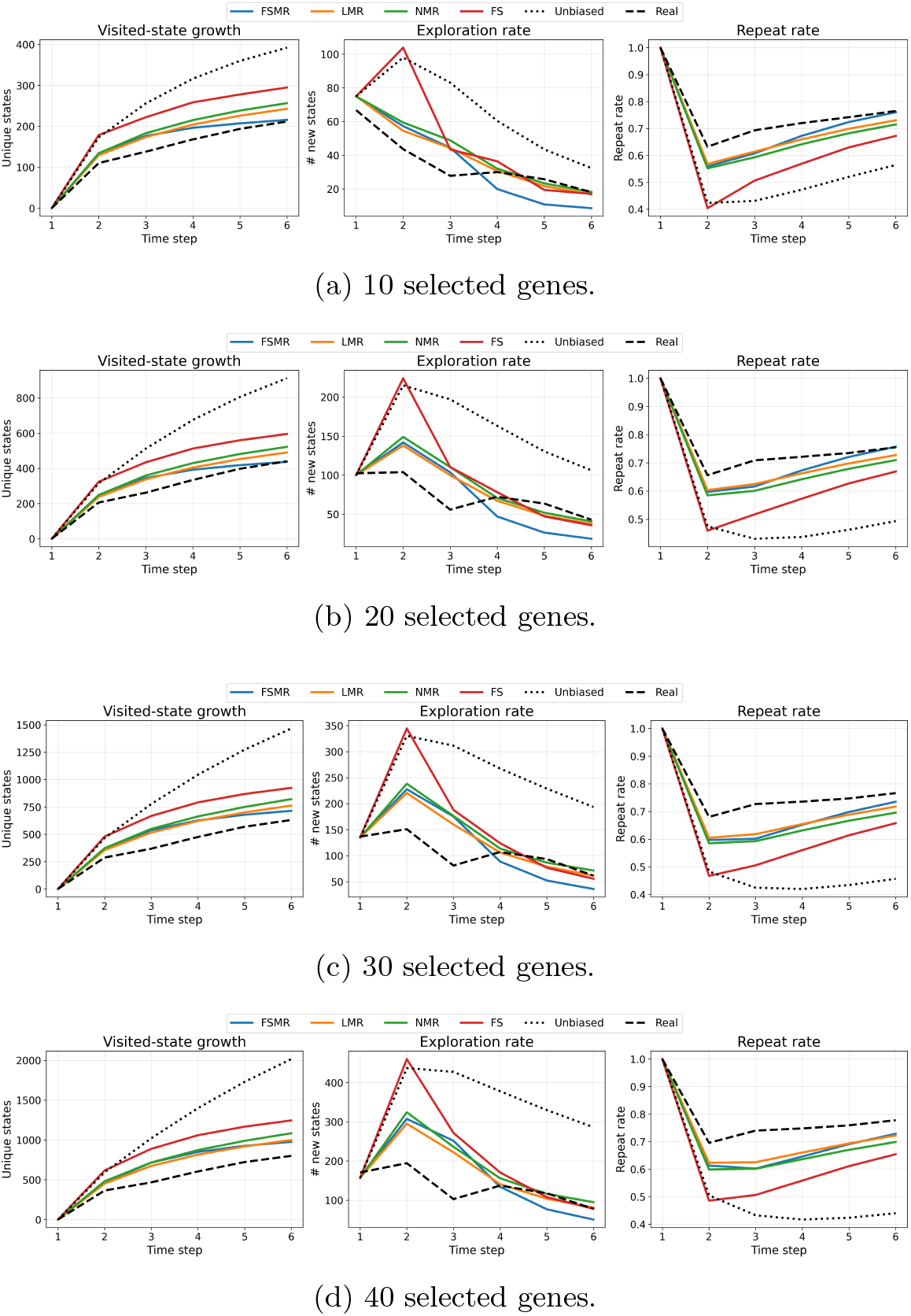
Dynamics comparison for dataset GSE120132 across different feature subset sizes.

**Fig. 4:**
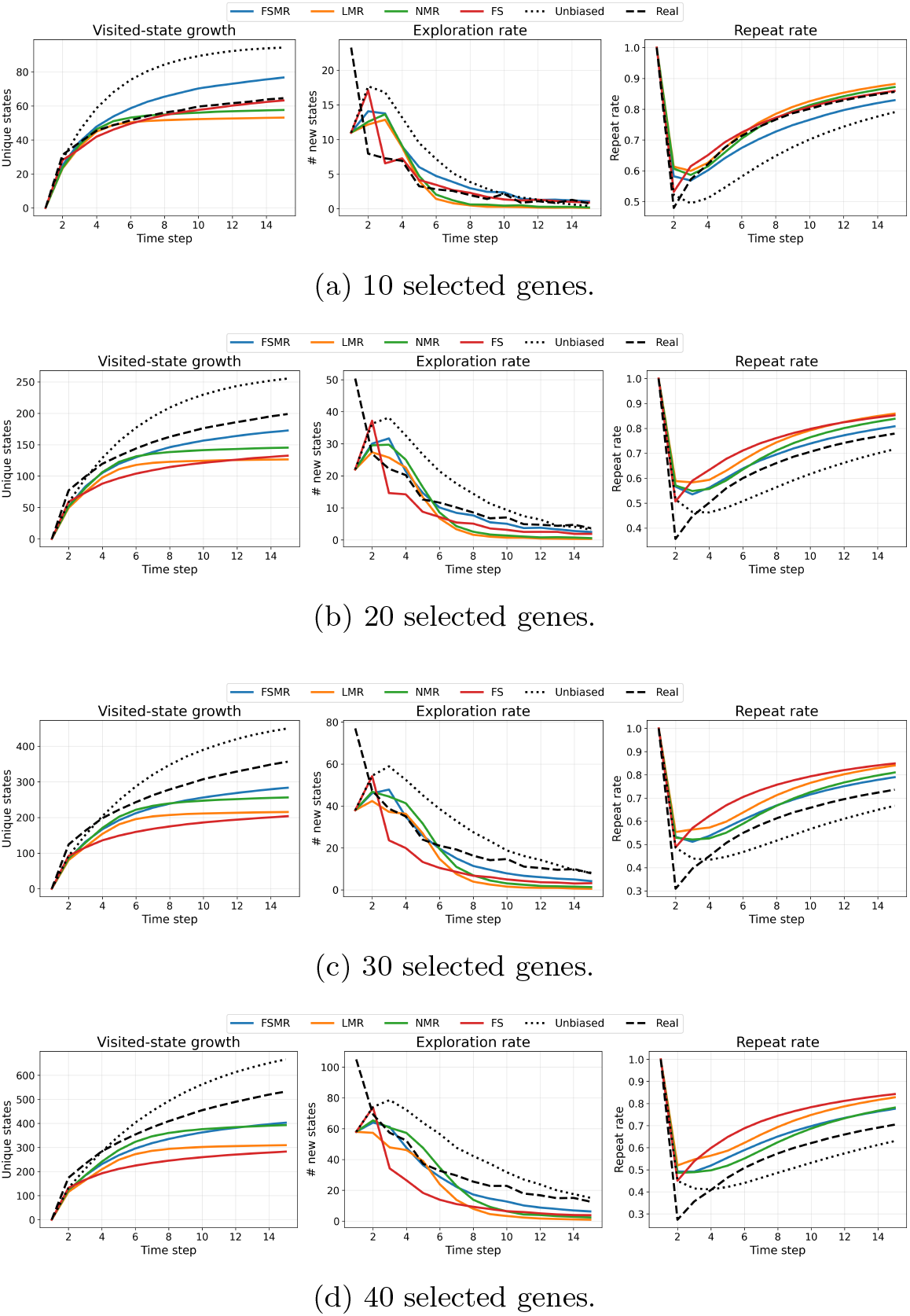
Dynamics comparison for dataset GSE164127 across different feature subset sizes.

Additionally, a one-sided paired t-test for the difference in means was performed across bootstrap runs to assess whether the biased models better reproduce the observed metrics than the unbiased model. For each bootstrap replicate, we computed the mean squared error (MSE) between the simulated and observed metric curves.

The null and alternative hypotheses were

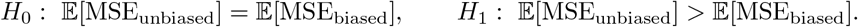

Here, a significant p-value indicates that the biased model has a statistically smaller MSE of the observed metric than the unbiased model. We perform this test for each bias model in Tab. 1 across 100 bootstrap runs. Results for each dataset and each set of selected genes are given in the tables 2–5.

**Table 2:** P-values for dataset GSE164127. Significant results are reported as *p <* 0.05.

| Metric | Model | 10 genes | 20 genes | 30 genes | 40 genes |
| --- | --- | --- | --- | --- | --- |
| Visited-state growth | FSMR | $p < 0.05$ | $p < 0.05$ | $p < 0.05$ | 0.087 |
| | LMR | $p < 0.05$ | 0.831 | 1 | 1 |
| | NMR | $p < 0.05$ | $p < 0.05$ | $p < 0.05$ | $p < 0.05$ |
| | FS | $p < 0.05$ | 0.972 | 1 | 1 |
| Exploration rate | FSMR | $p < 0.05$ | $p < 0.05$ | $p < 0.05$ | $p < 0.05$ |
| | LMR | $p < 0.05$ | $p < 0.05$ | $p < 0.05$ | $p < 0.05$ |
| | NMR | $p < 0.05$ | $p < 0.05$ | $p < 0.05$ | $p < 0.05$ |
| | FS | $p < 0.05$ | $p < 0.05$ | $p < 0.05$ | 0.778 |
| Repeat rate | FSMR | $p < 0.05$ | $p < 0.05$ | 0.257 | 0.997 |
| | LMR | $p < 0.05$ | 0.999 | 1 | 1 |
| | NMR | $p < 0.05$ | 0.117 | 0.427 | 0.745 |
| | FS | $p < 0.05$ | 1 | 1 | 1 |

**Table 3:**
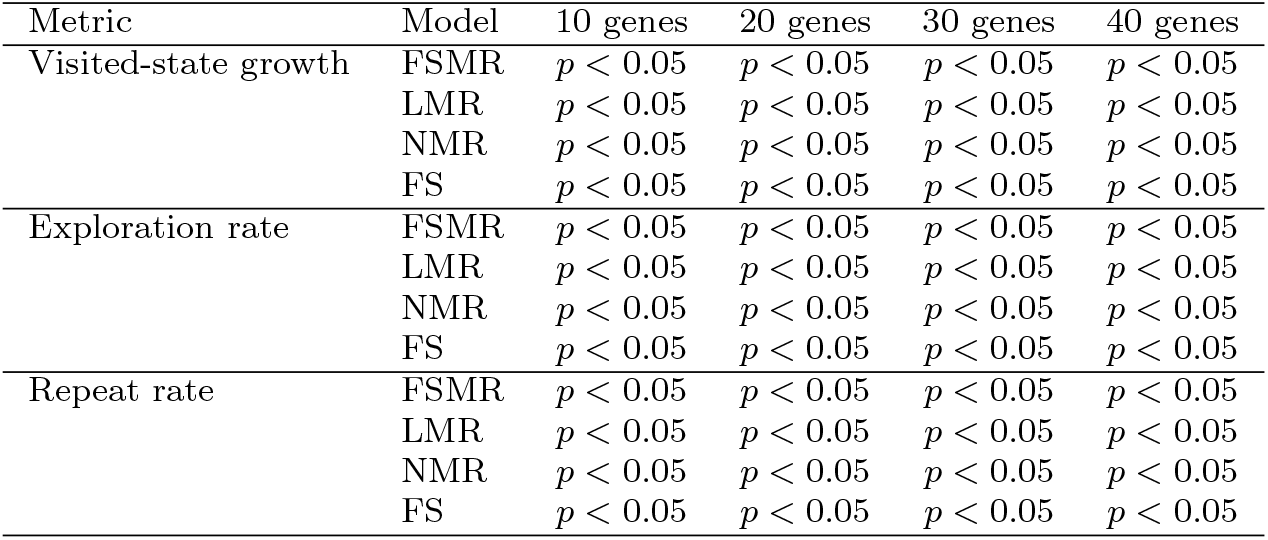
P-values for dataset GSE120132. Significant results are reported as *p <* 0.05.

**Table 4:**
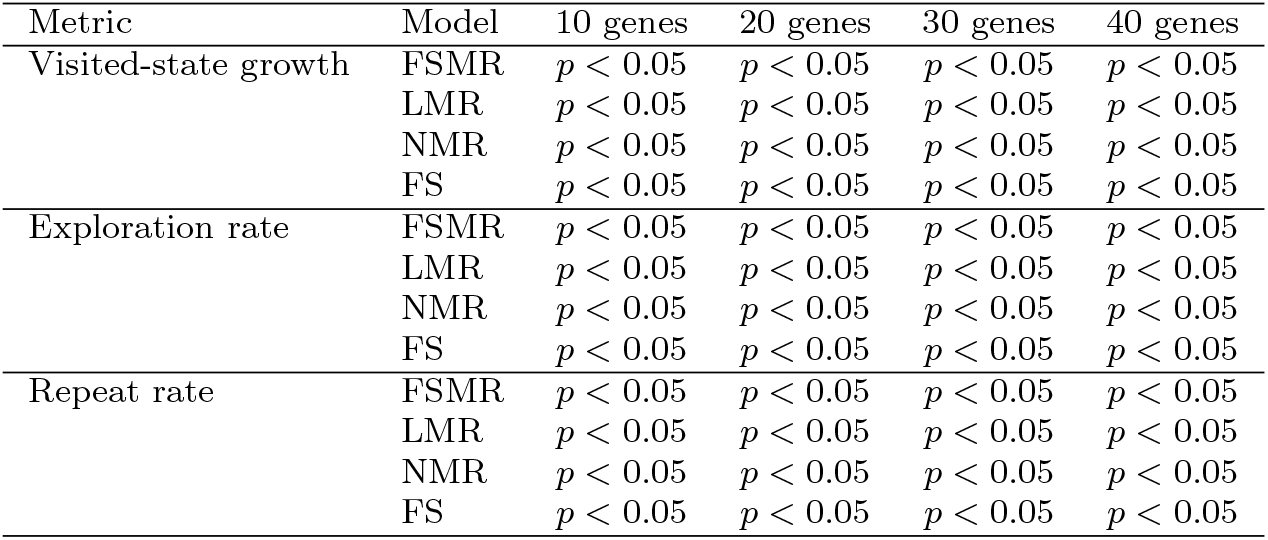
P-values for dataset GSE161141. Significant results are reported as *p <* 0.05.

**Table 5:**
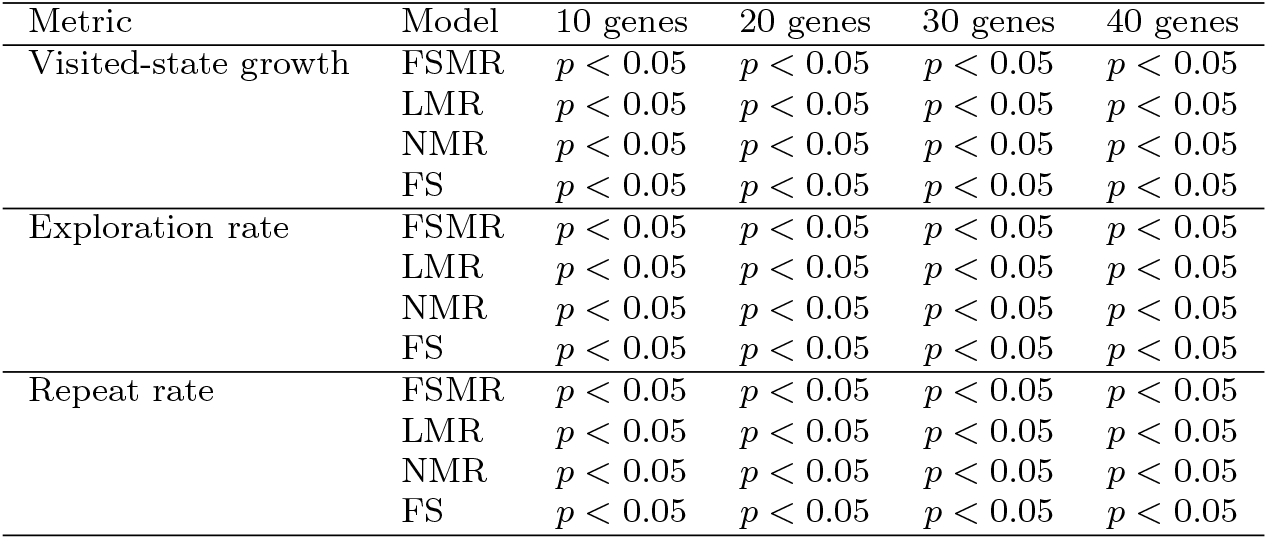
P-values for dataset GSE80672. Significant results are reported as *p <* 0.05.

**Table 6:**
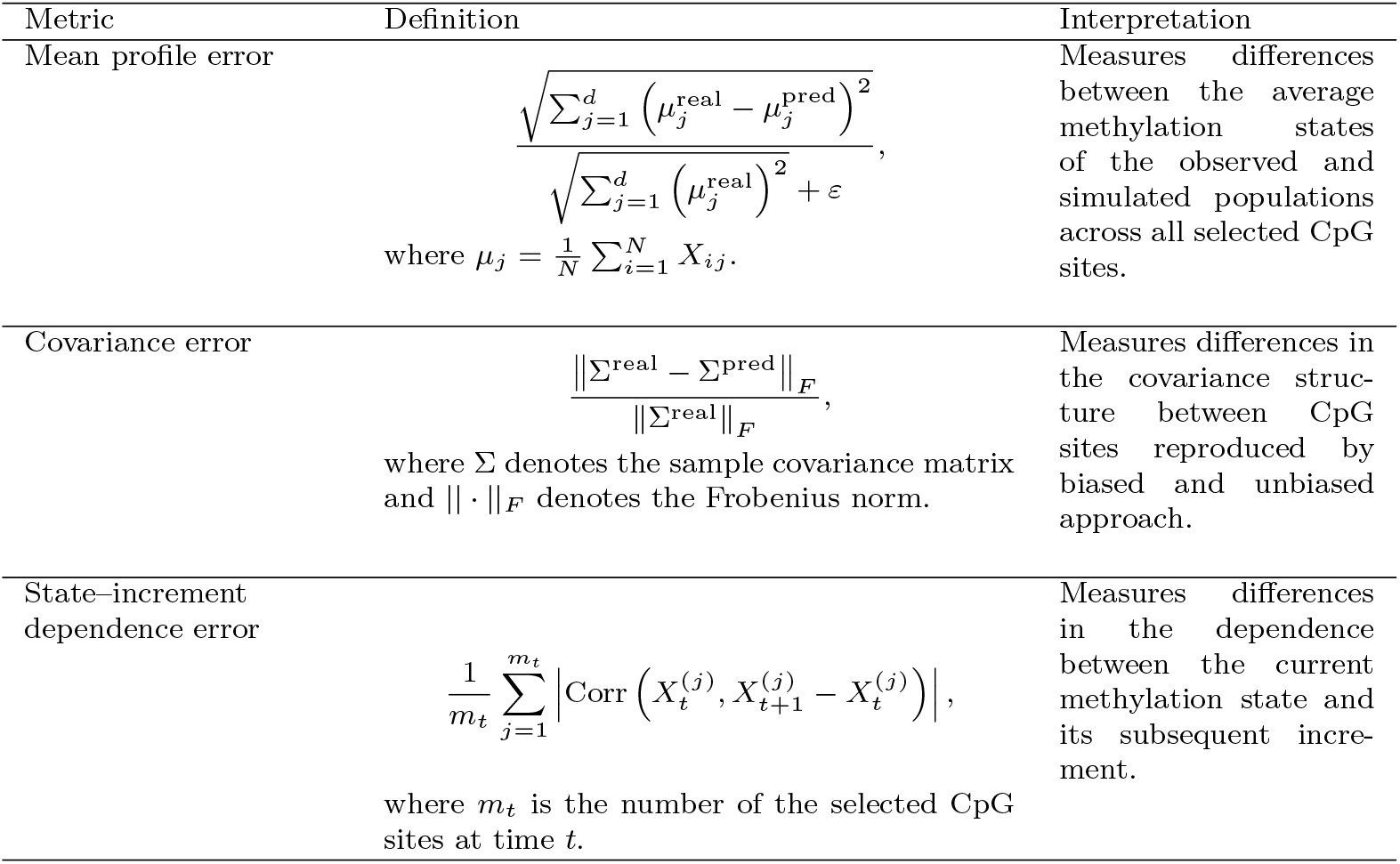
Performance metrics used to evaluate the predicted methylation trajectories. Lower values indicate better agreement.

## 3 Transition kernel estimation

The analysis in the previous section focused on the exploratory properties of biased and unbiased processes. The results presented in Fig. 1–3 and Tab. 4–2 show that processes with a regulatory bias reproduce the observed patterns of methylation state-space exploration more closely than purely stochastic models. This suggests that state-dependent effects indeed influence aging dynamics. We therefore next investigate whether accounting for such effects improves the prediction of future methylation states. To do so, we compare two approaches. The first learns a state-dependent bias directly from the data and uses the resulting model for prediction. The second assumes no regulatory bias and predicts future states using only the stochastic component of the dynamics. We hypothesize that incorporating a learned bias will lead to more accurate predictions. The next step in our study is therefore to empirically learn the form of the bias term from data, followed by the prediction of future system states based on the inferred model.

As Markov chains, the processes (1) and (2) are completely defined by their initial state and their transition probability kernel. Intuitively, the transition kernel describes how the system evolves from one state to the next: given the current state *X*_*t*_, it specifies the probability distribution of the next state *X*_*t*+1_. In our case, both processes (2) and (1) have continuous state space, which means the transitions are defined in terms of *regions* rather than states. Formally, the transition kernel is defined as

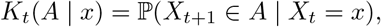

where *A* is a region (i.e., a subset of the state space). This describes the probability that the process moves from the current state *x* to region *A* in one step. The knowledge of these probabilities is essential to understand the mechanism of movement and changes within the process, and also to predict the process dynamic in the future time points that are not yet observed. Hence, the next step of our study is to empirically estimate the transition probabilities for both the biased and unbiased processes using data from Section 2.1 again. We then use the estimated kernel to generate the process dynamic further and evaluate its accuracy.

Given observed trajectories 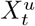 and 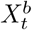, we estimate the transition kernel from consecutive observations (*X*_*t*_, *X*_*t*+1_). The aim is to learn how the methylation state changes from one time point to the next and to use this information to predict future states. Since both the biased and unbiased models assume Gaussian random noise in the update equations (1)–(2), the resulting transition kernels are naturally described by multivariate normal distributions. In other words, the probability of transitioning from one state to another can be fully characterized by a multidimensional Gaussian distribution whose shape is determined by its mean vector and covariance matrix. To enable the generation of new states in both processes, it is therefore necessary to estimate their mean and covariance parameters from data. To predict the future state, we will look at the process increments. For the unbiased process (1), the increments satisfy

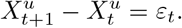

In this simple case, the increments have the same distribution as *ϵ*_*t*_. Under this assumption, no regulation in the dynamics is imposed, and consequently there is no structure to learn from the data. It remains to only estimate the covariance matrix. To do so, we use the Ledoit–Wolf shrinkage estimator [25], which is normally used when the number of observations is small relative to the number of genes. The covariance matrix is then estimated from the observed increments

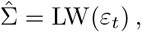

where LW(·) denotes the Ledoit–Wolf shrinkage estimator. The transition to the next state, given the current state *x*, is defined as follows

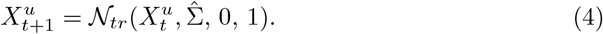

This means that the next state is obtained by adding random fluctuations to the current state, with variability estimated directly from the data. The future state is obtained solely from the current state and an independent random perturbation.

For the biased process (2), the presence of the bias term *b*(*X*_*t*_) introduces additional structure into the dynamics. At this point, we hypothesize that real methylation dynamics contain an underlying structure that is not captured by independent noise alone. By learning this structure directly from the data through the bias term, we expect to obtain a more accurate description of methylation trajectories than with the unbiased approach (4). Then, in addition to the current state of the process and normal noise, we need to learn the bias term from data. To do so, we model the *increments*

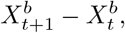

which represents the change of the process between two consecutive time points. Using the observed pairs 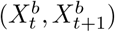, we estimate the conditional mean drift

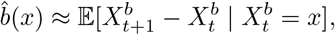

that is, the average change of the process conditioned on the current state *x*. In other words, 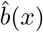 describes the expected direction and magnitude of movement of the system at state *x*. In subsequent experiments, we estimate 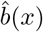 using linear regression with elastic net regularization. Specifically, we approximate the drift by

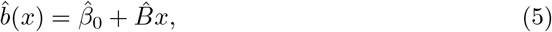

where 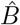 is the estimated coefficient matrix and 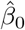 is the intercept vector. where the coefficients are learned from the observed transitions. Note that our goal is not to obtain the most accurate predictor, but rather to demonstrate that introducing a drift component improves the description of the dynamics compared to a purely unbiased process; for this reason, we intentionally adopt the rather simple linear regression model. Once the drift is estimated, the residuals are computed as

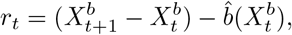

which is a stochastic component not explained by the learned drift. Similarly to the unbiased approach, we estimate the covariance matrix from the residuals using the LW estimator

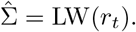

The resulting learned transition model is therefore given by

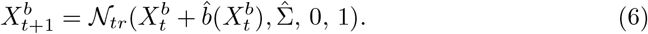

Thus, each update step uses an estimated deterministic drift 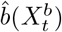. Future trajectories are sampled iteratively by repeatedly applying this learned transition rule.

To validate the two approaches, we perform the following experiment. We use the same datasets as before (GSE164127, GSE120132, GSE161141, and GSE80672), each consisting of trajectories 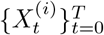 for multiple individuals *i*. Since we want to estimate the future trajectories, the data is split chronologically along the time axis, with the first 80% of the time points used for training and the remaining 20% used for validation. Using the training portion, we learn the new states using both approaches and (6). In the former case, we estimate the transition kernel simply using the current state as a mean and an additional noise term. In the biased approach, we learn the process slope term using elastic net regression. Then, starting from the last observed states in the train set, we simulate forward trajectories using unbiased (4) and biased (6) approaches. These generated trajectories are then compared to the true observed trajectories for validation.

### 3.1 Performance evaluation

The proposed model is evaluated by comparing the simulated trajectories with the test data using three complementary metrics: differences in the mean, covariance, and state–increment dependence. The first two metrics quantify the agreement between the marginal distributions of the simulated and observed methylation states, whereas the third evaluates whether the simulated process reproduces the dependence between the current methylation state and its increment. Recall that our objective is to capture the shape of drift and direction of methylation changes, rather than to exactly reconstruct individual methylation states, for which these three metrics appear to be the most informative. Since the model is stochastic, the futures states are simulated independently *n* = 100 times. The corresponding metrics are computed for each simulation and averaged across all runs. Finally, the average values over the prediction horizon are reported for each bootstrap sample. The exact formulas and description of the metrics are given in Table. 6. The final results are shown in Fig. 5–8 and Fig. 10.

**Fig. 5:**
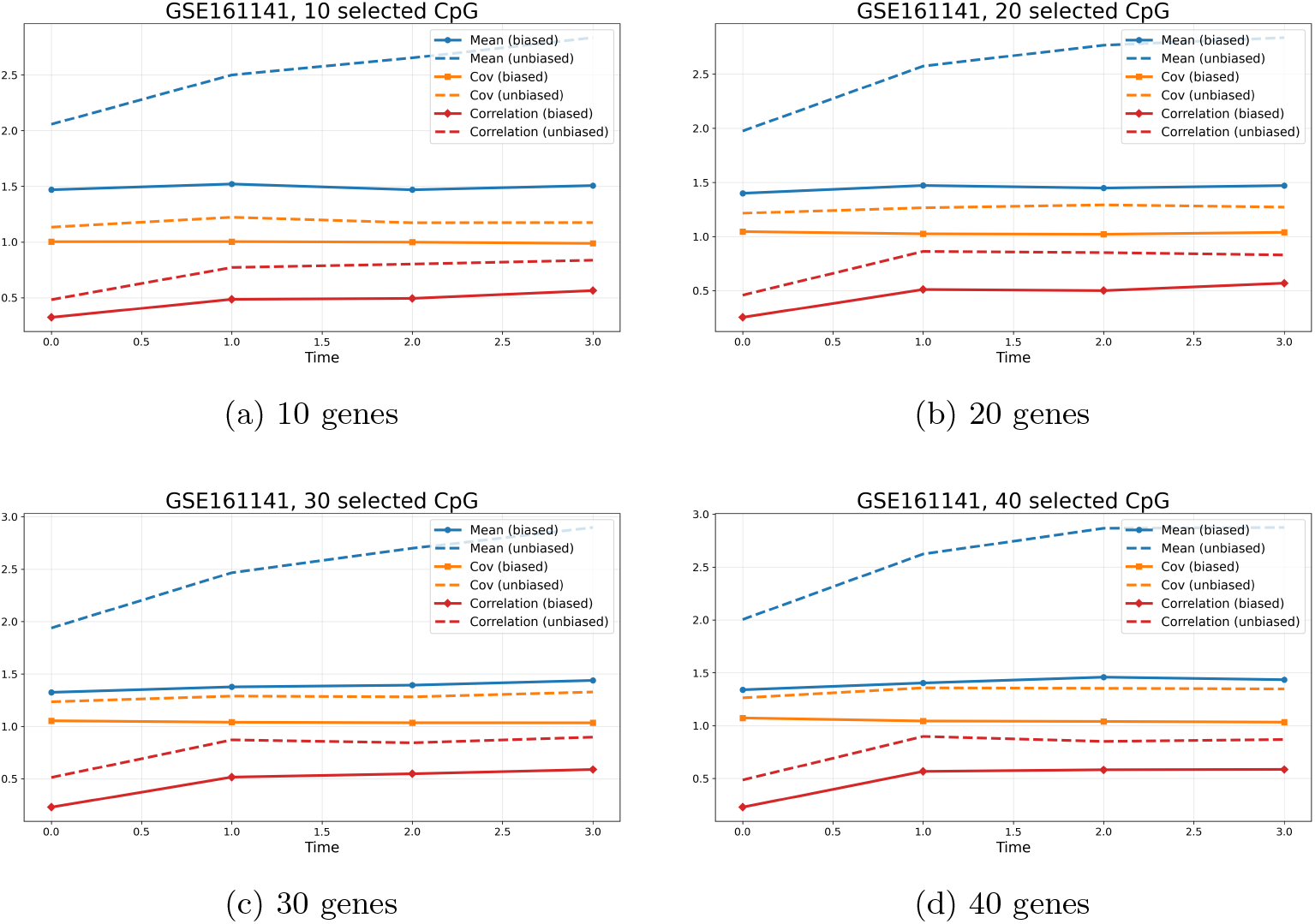
Prediction errors for biased and unbiased dynamics for dataset GSE161141.

**Fig. 6:**
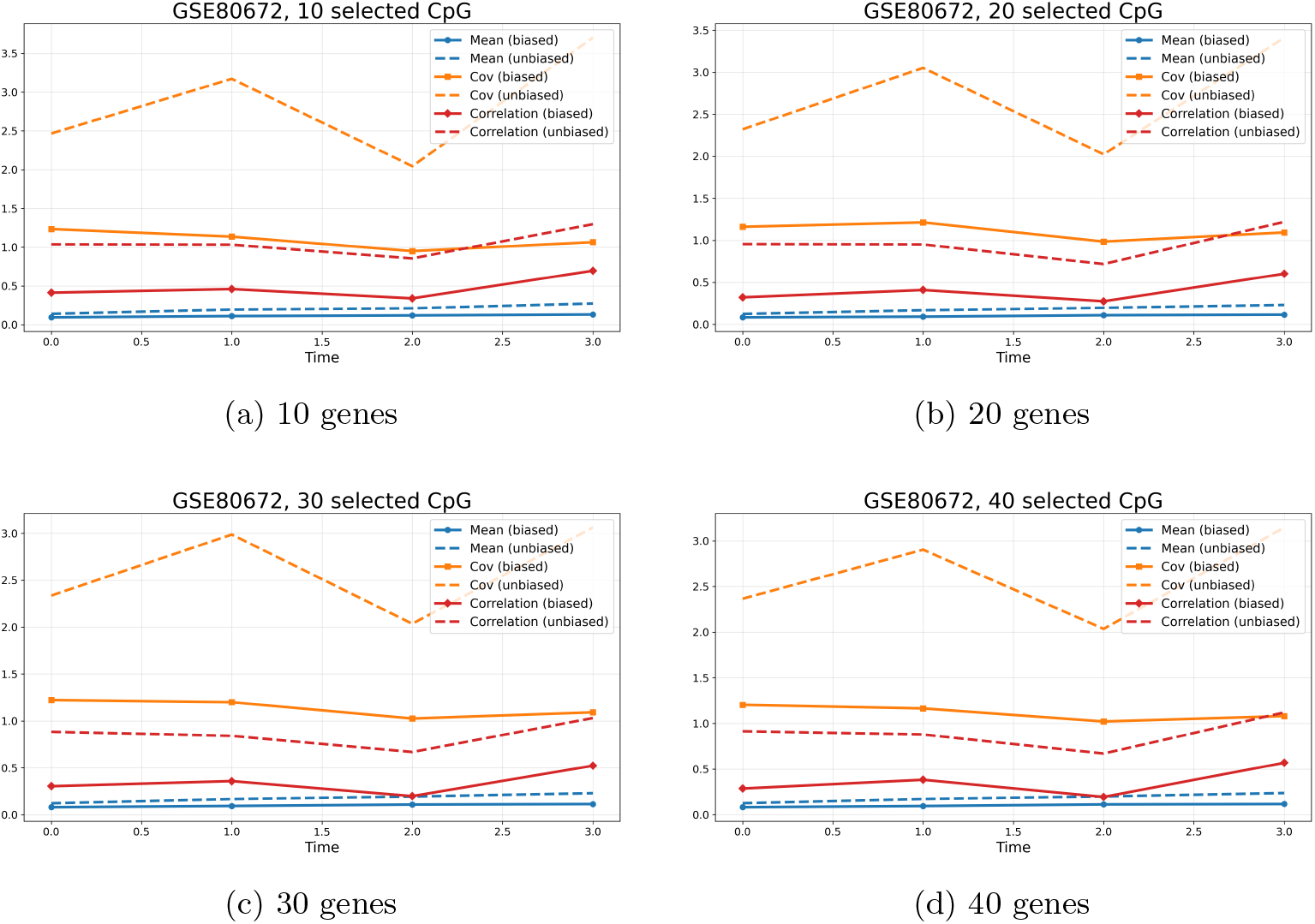
Prediction errors for biased and unbiased dynamics for dataset GSE80672.

**Fig. 7:**
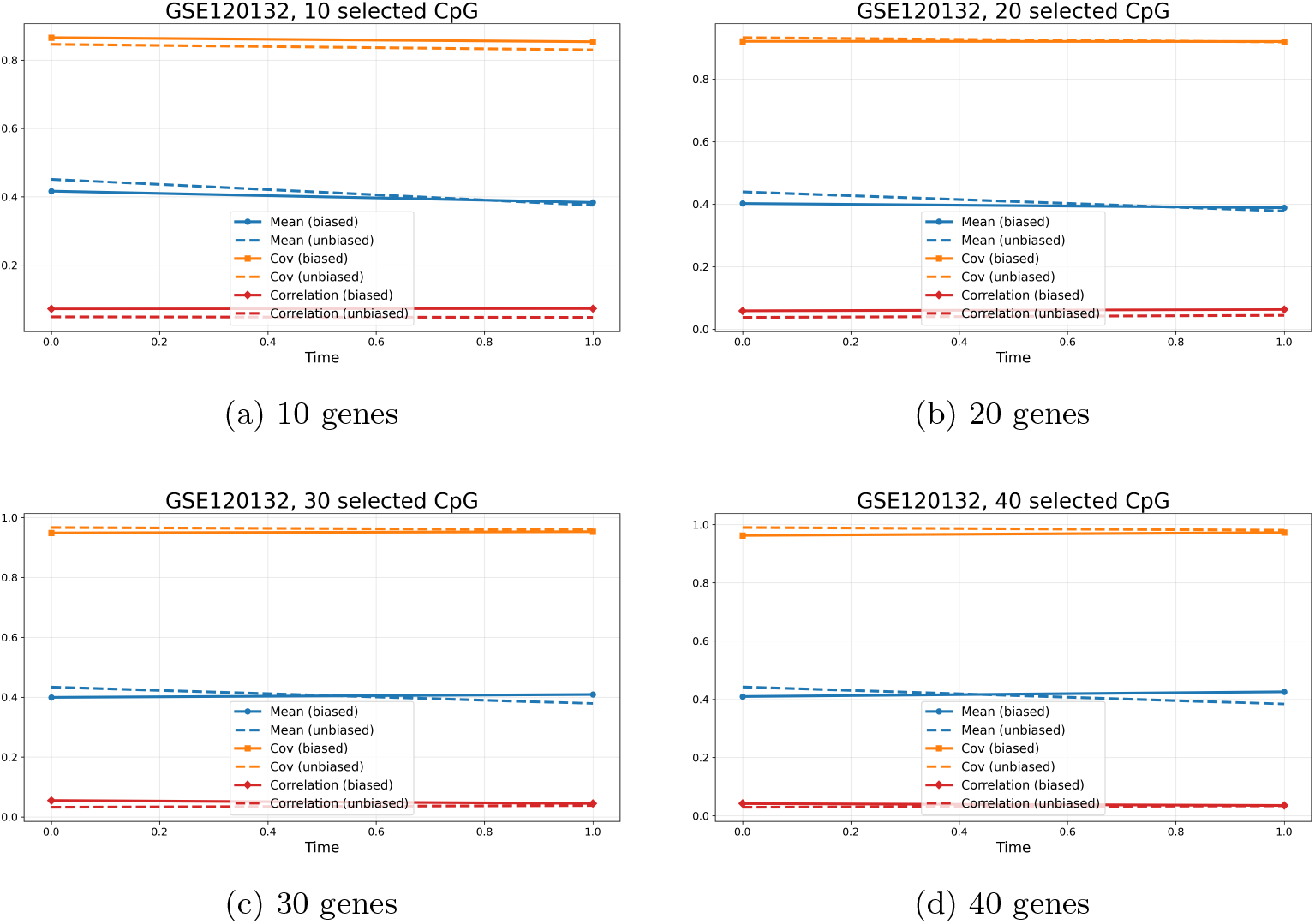
Prediction errors for biased and unbiased dynamics for dataset GSE120132.

**Fig. 8:**
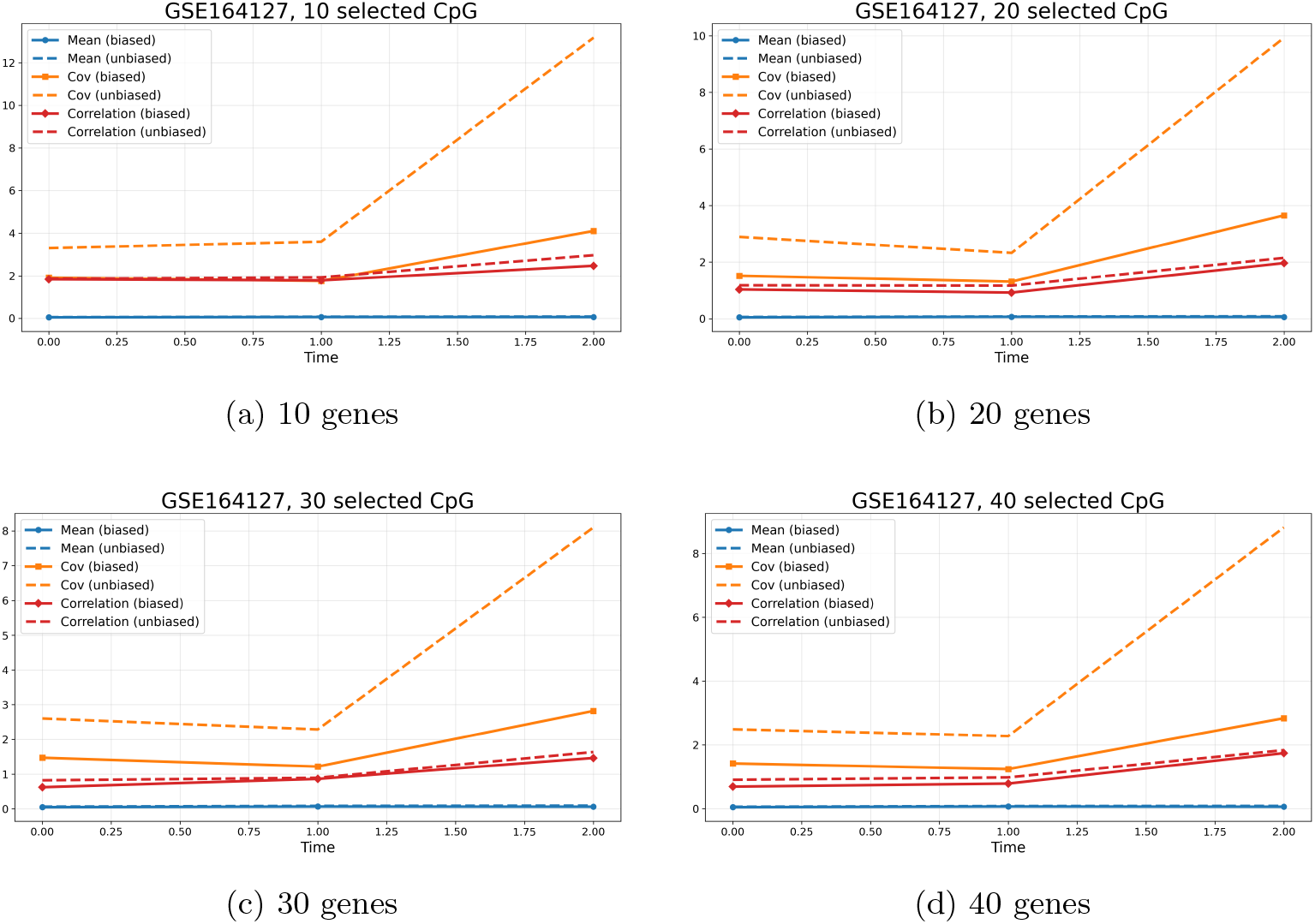
Prediction errors for biased and unbiased dynamics for dataset GSE164127.

**Fig. 9:**
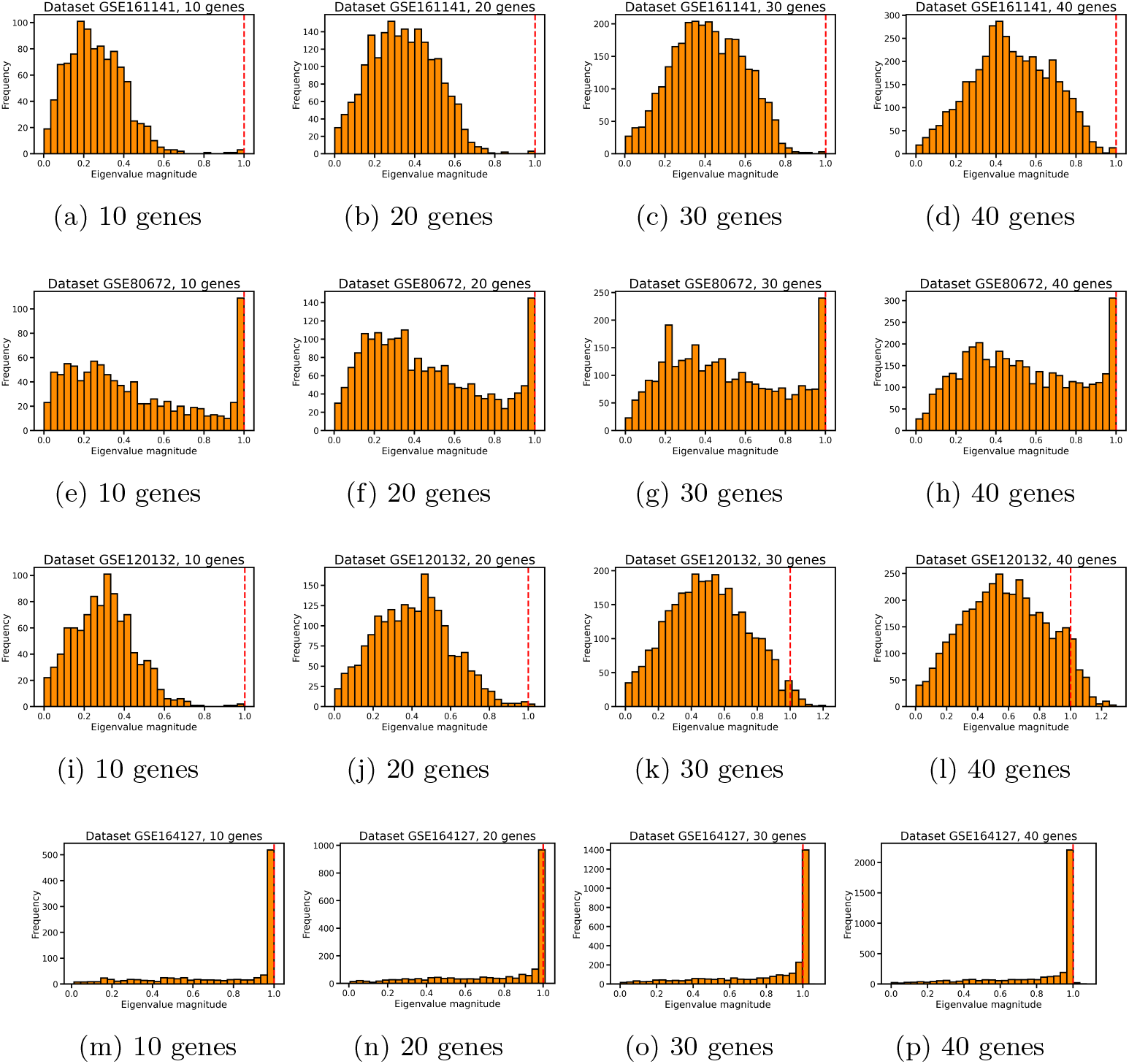
Eigen values | *λ* | across bootstrap runs for various datasets and subsets of selected genes.

**Fig. 10:**
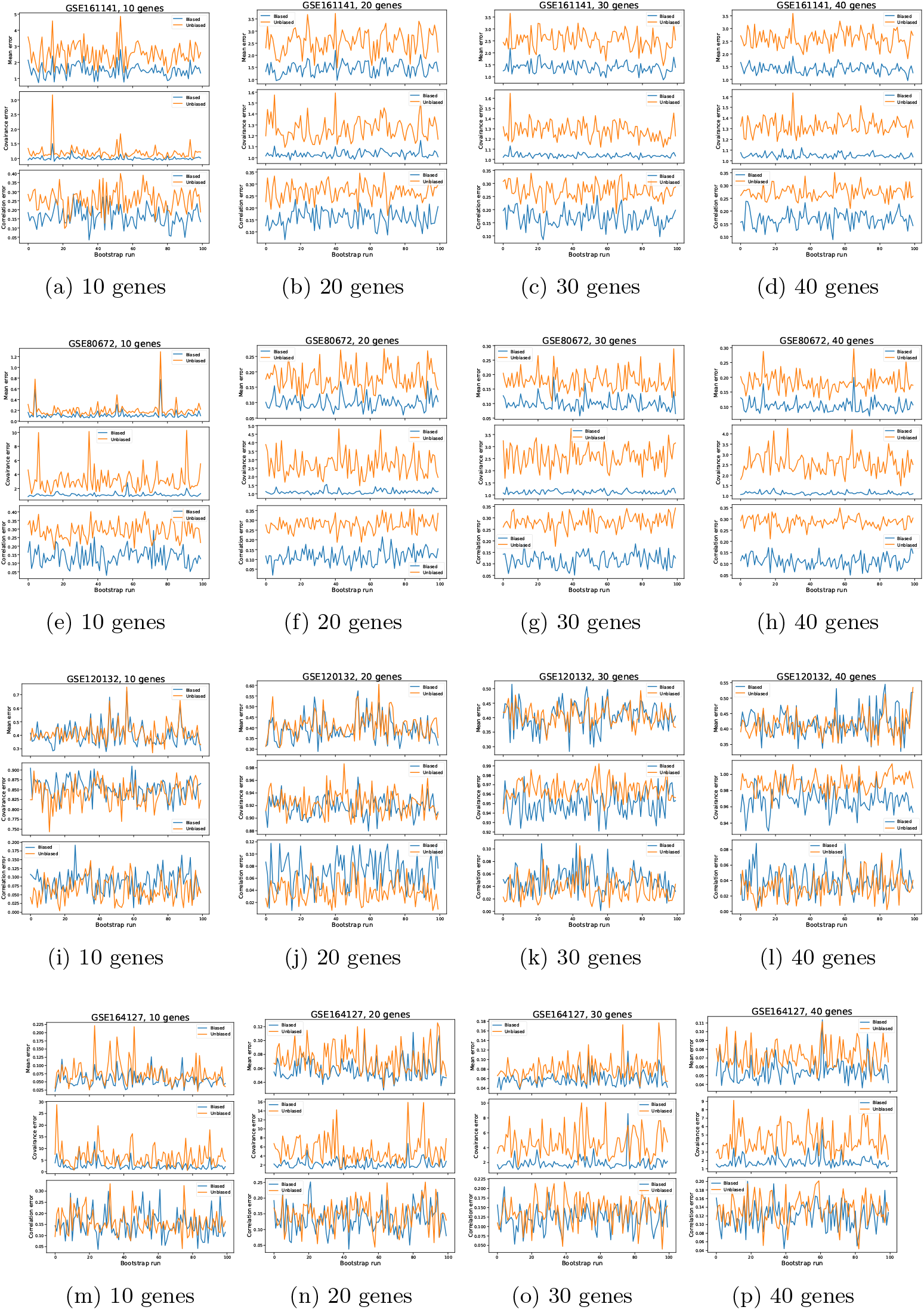
Prediction errors averaged over time points as a function of bootstrap replicate for the biased and unbiased models across all datasets and gene-selection settings.

As before, we perform the one-sided paired t-test to see if the average errors in biased and unbiased predictions differ significantly. We average all three metrics across time points, and perform the t-test on the bootstrap samples. The alternative hypothesis in this case is that average errors in the unbiased approach are larger than those in the biased approach. The results are given in Tab. 7–10.

To further investigate the contraction of the accessible state space, we analyzed the eigenvalues of the estimated update matrix

**Table 7:** P-values for prediction error metrics for dataset GSE164127. Significant results are reported as *p <* 0.05.

| Metric | 10 genes | 20 genes | 30 genes | 40 genes |
| --- | --- | --- | --- | --- |
| Mean error | $p < 0.05$ | $p < 0.05$ | $p < 0.05$ | $p < 0.05$ |
| Covariance error | $p < 0.05$ | $p < 0.05$ | $p < 0.05$ | $p < 0.05$ |
| State-increment dependence error | $p < 0.05$ | $p < 0.05$ | $p < 0.05$ | $p < 0.05$ |

**Table 8:** P-values for prediction error metrics for dataset GSE120132. Significant results are reported as *p <* 0.05.

| Metric | 10 genes | 20 genes | 30 genes | 40 genes |
| --- | --- | --- | --- | --- |
| Mean error | $p < 0.05$ | $p < 0.05$ | 0.304 | 0.874 |
| Covariance error | 1 | $p < 0.05$ | $p < 0.05$ | $p < 0.05$ |
| State-increment dependence error | 1 | 1 | 1 | 0.999 |

**Table 9:**
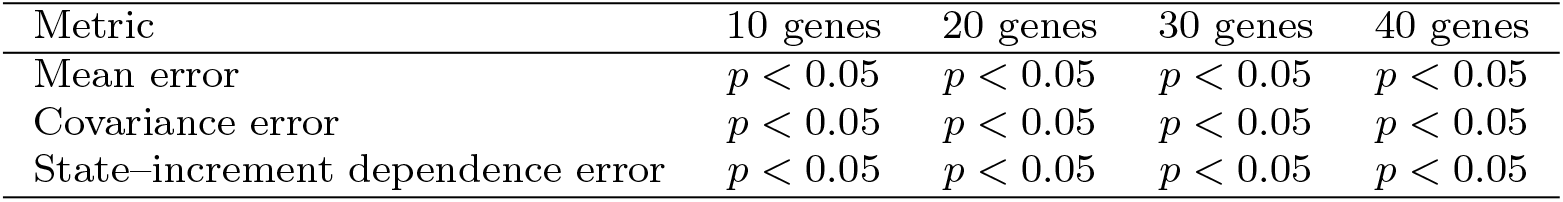
P-values for prediction error metrics for dataset GSE161141. Significant results are reported as *p <* 0.05.

**Table 10:**
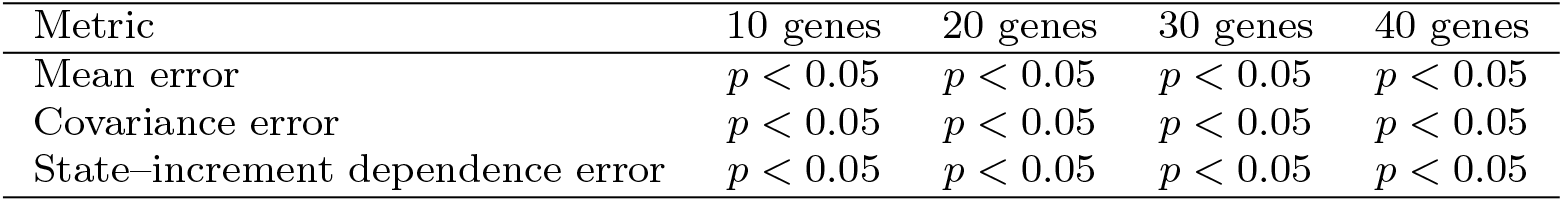
P-values for prediction error metrics for dataset GSE80672. Significant results are reported as *p <* 0.05.

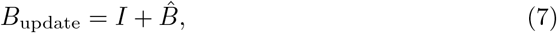

Where 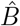 is the estimated drift matrix from (5) and *I* is the identity matrix. The eigenvalues were computed by solving

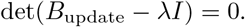

Eigenvalues with | *λ <* 1 | correspond to contractive directions, whereas | *λ >* 1 | indicate expansion. Consistent with previous experiments, eigenvalues were computed for 100 bootstrap replicates and their distributions were compared. The box plots of the resulted eigenvalues over bootstrap runs are given in Fig. 9

## 4 Discussion

Fig. 1–4 illustrates the effect of introducing various forms of regulatory bias into the dynamics. The plots compare the biased process (2) with various forms of drift term (Tab. 1) against unbiased (1) and empirical (real) processes across multiple datasets and different sets of selected genes. The results in Tab. 4–5 report the significance results of the comparatives. The most visible difference appears in the visited-state growth. The unbiased process rapidly explores the state space and continuously visits a large number of previously unseen states. In contrast, the biased processes explore the state space more slowly and reach a substantially smaller number of unique states over time. The empirical process is even more constrained and typically shows the slowest state-space expansion. This effect becomes especially visible in the GSE161141 data, where the difference between the unbiased and empirical trajectories increases steadily over time. The exploration-rate curves confirm this behavior. The unbiased process maintains a relatively high rate of discovering new states, particularly during the early stages of the simulation. By comparison, the biased processes quickly reduce their exploration activity and increasingly revisit already observed configurations. The empirical process often exhibits the strongest decline in exploration rate, especially in the GSE161141 data, where only a small number of new states continue to appear after several iterations. These results suggest that the real methylation dynamics evolve within a restricted subset of the accessible state space. The repeat-rate diagnostics further support this interpretation. In the empirical process, the repeat rate rapidly approaches one, indicating that trajectories repeatedly return to previously visited states instead of continuously exploring new configurations. The biased processes show a similar tendency, although less strongly, while the unbiased process maintains significantly lower repeat rates throughout the simulation. These observations are supported by one-sided paired t-tests across bootstrap replicates, which showed that, for the absolute majority of datasets, gene subsets, and exploration metrics, the biased models produced significantly smaller discrepancies from the empirical dynamics than the unbiased model It is worth mentioning that while the different bias formulations produce slightly different trajectories, they all show the same overall behavior. The same applies to various subsets of genes. In all datasets, regardless of the number of selected genes, introducing a bias reduces the growth of the visited state space, decreases the rate of discovering new states, and increases the tendency to revisit previously observed states. Some bias models match the empirical dynamics more closely for certain datasets, while others perform better in different settings, suggesting that the exact form of the bias is dataset-dependent. However, the presence of a bias, regardless of its form, consistently moves the dynamics closer to the empirical process, whereas the unbiased model systematically overestimates exploration. Importantly, the plots demonstrate that the empirical methylation dynamics show an even stronger restriction of the accessible state space than any of the simulated models. This means that, as aging progresses, the set of accessible methylation states becomes increasingly restricted. The experiments show that the purely noise-driven process fails to capture this progressive restriction and instead continues to explore new states. These results suggest that state-space restriction is an important characteristic of epigenetic aging that should be taken into account when defining and modeling the aging process.

Fig.5–8 evaluate the accuracy of the predicted process states obtained using the unbiased (4) and biased (6) approaches. Several complementary metrics are considered to evaluate both local and global properties of the simulated dynamics: mean error, covariance, and State–increment dependence error. The errors are computed by comparing the predicted states generated by the biased and unbiased approaches with the real dynamics. In the plots, solid lines correspond to the errors of the biased predictions relative to the real process, while dashed lines correspond to the errors of the unbiased predictions relative to the real process. The mean error measures the difference between the average feature values of the predicted and empirical states. The covariance error measures how well the joint variability between features is preserved, whereas the state–increment dependence error measures how accurately the dependence between the current state and its subsequent change is captured. Across most datasets, the biased approach consistently produces smaller errors than the unbiased one, which is mostly visible in mean and covariance errors. This indicates that the biased dynamics preserve the structure of the empirical data more accurately. The difference is particularly visible in datasets GSE80672 and GSE161141, where the unbiased process diverges progressively from the empirical trajectories over time. These conclusions are further supported by the statistical test results given in Tab. 7 –10. The results indicate that errors in the biased approach are significantly smaller than the errors in unbiased approach for most of the metrics and dataset. The exception is error in dataset GSE120132, which shows no significant difference from the unbiased approach. However, as can be seen in Fig. 10, in this case, state–increment dependence errors in both biased and unbiased predictions are almost the same, which indicates possible lack of correlation between genes. Moreover, the are only 6 available time points in the dataset, whereof only 2 were used for validation. Hence, a possible explanation of a non-significant result in state–increment dependence error is lack of train data and validation data. Fig. 9 shows the distribution of the eigenvalues | *λ* | of the learned update *B*_*update*_ in (7) across bootstrap replicates. In all datasets, most eigenvalue magnitudes are below one, which indicates that the learned dynamics are contractive in most directions. The strongest contraction is observed for the GSE161141 and GSE120132 datasets, where most eigenvalues are concentrated well below one. In contrast, GSE164127 contains many eigenvalues close to one, suggesting that a a large number of dimensions in the dynamics is nearly neutral, while contraction occurs only in a subset of directions. This is consistent with the exploration results shown in Fig. 4, where the exploration rate in empirical process exceeds that of all biased models and becomes closer to the unbiased behavior. Nevertheless, both analyses indicate the presence of state-space contraction, although the effect is substantially weaker than in the other datasets. This observation is also biologically plausible, as some CpG sites remain relatively stable during aging, whereas others change more substantially. Therefore, the learned dynamics are not expected to be contractive in all directions (i.e. that all eigenvalues are below 1). Instead, the eigenvalue spectrum suggests that contraction occurs only in a subset of dynamical modes, while the remaining modes are approximately neutral.

## 5 Conclusion

Overall, the results show that introducing a regulatory bias significantly changes the qualitative behavior of the process. The unbiased dynamics over-explore the state space and produce increasingly unrealistic variability. In contrast, the biased process generates more restricted trajectories and remains consistently closer to the empirical data. These findings support the interpretation that aging-related dynamics are not well described by pure noise stochastic exploration, but instead evolve under increasing structural restrictions that eventually limit the accessibility of states over time. Moreover, the observed pattern of state-space shrinkage is consistent with discrete bifurcation events, suggesting that resilience declines through a series of transitions. At each bifurcation, part of the state space becomes inaccessible, reducing the number of trajectories available to the system. As these losses accumulate, future dynamics become increasingly constrained and the range of possible interventions narrows. In this view, the timing of interventions becomes critical, since opportunities to redirect the system may be lost once a bifurcation has occurred.

## Methods

The aim of the study was to determine whether incorporating regulatory bias improves the description and prediction of epigenetic aging dynamics compared with a purely noise-driven stochastic process. The setting consisted of publicly available DNA methylation datasets from mice, rats, and bats (GSE120132, GSE164127, GSE161141, and GSE80672). DNA methylation measurements were organized into tensors of dimension time *×* observations *×* features. For state-space analysis, methylation values were discretized using an adaptive bin width Δ = *d*^*™*3*/*2^, with *d ∈ {*10, 20, 30, 40*}* selected CpG sites. Robustness was assessed using 100 bootstrap replicates per feature-set size (400 datasets per GEO dataset). Biased and unbiased dynamics were compared through visited-state growth, exploration rate, and repeat rate. State-dependent drift was estimated using elastic-net regression (*α* = 10^*™*3^, *l*_1_ ratio = 0.5, 10^4^ maximum iterations), and covariance matrices were estimated using the Ledoit–Wolf shrinkage estimator. Models were trained using an 80/20 train–test split and evaluated over 100 stochastic simulations per bootstrap. Statistical significance was assessed using one-sided paired t-tests across bootstrap replicates, testing whether the biased model yielded lower MSE values and lower prediction errors (mean, covariance, and state– increment dependence errors) than the unbiased model. In addition, the eigenvalues of the estimated matrix update were computed across all bootstrap replicates to assess the degree of contraction of the accessible state space.

### Abbreviations

*AR(1)*: First-order autoregressive model
*CpG*: Cytosine-phosphate-guanine dinucleotide site
*GEO*: Gene Expression Omnibus
*LW*: Ledoit–Wolf shrinkage estimator
*FSMR*: Feature-specific regulation with mean reversion
*FS*: Feature-specific regulation
*LMR*: Linear mean reversion
*NMR*: Nonlinear mean reversion

## Declarations

### Ethics approval and consent to participate

This study used exclusively publicly available, de-identified DNA methylation datasets obtained from the Gene Expression Omnibus (GEO). No new human or animal subjects were recruited, and the authors performed no experimental procedures. Ethics approval and consent were therefore not required.

### Consent for publication

Not applicable.

### Availability of data and materials

All datasets analyzed during this study are publicly available through the Gene Expression Omnibus (GEO) repository under accession numbers GSE120132, GSE164127, GSE161141, and GSE80672. Additional code and processed data will be made available by the corresponding author upon request.

### Authors’ contributions

All authors contributed equally to the manuscript.

